# STING activation of IRF3 is tuned by PELI2 to suppress basal activation and reinforce the anti-viral response

**DOI:** 10.1101/2023.04.15.537029

**Authors:** Christopher Ritchie, Lingyin Li

## Abstract

The STING pathway is the first line of defense against a variety of threats. STING activation leads to two main signaling branches, IRF3 signaling and NF-κB signaling, that have differential roles in anti-cancer, anti-viral, and anti-bacterial immunity and autoimmunity. However, it is unknown how these two signaling branches are differentially regulated depending on context. Here, we identify PELI2 as a regulator of STING that preferentially inhibits IRF3 signaling while enhancing NF-κB signaling. Mechanistically, we show that PELI2 inhibits IRF3 signaling by binding to phosphorylated Thr354 and Thr356 on the C-terminal tail of STING, leading to ubiquitination and function switching of TBK1. PELI2 is expressed under basal conditions to suppress IRF3 signaling and prevent interferonopathies. During viral infection, however, STING signaling rapidly downregulates PELI2 to unleash production of anti-viral type-I interferons. Normally, PELI2 levels are restored following viral clearance. However, lupus patients have insufficient PELI2 levels and high basal interferon production, suggesting dysregulation of PELI2 may have a causative role in lupus and other interferonopathies.

## Introduction

The human body constantly encounters threats on a cellular level that it must properly respond to in order to survive. These threats originate both externally, such as viral and bacterial infection, as well as internally, such as the development of cancer. The first step in properly responding to these threats is detection. Human cells are equipped with an array of innate immune pathways that detect threats through a variety of danger signals. One pathway that is nearly ubiquitous in the immune response is the cGAMP-STING pathway, which detects mislocalized double stranded DNA (dsDNA), a danger signal associated with many threats, including pathogen infection, cellular damage, and cancer. Following detection of mislocalized dsDNA, the enzyme cGAS is activated and synthesizes the small molecule second messenger of the pathway, 2’3’-cyclic-GMP-AMP (cGAMP), which serves as an agonist for the pathway’s receptor, STING. Once activated by cGAMP, STING can trigger multiple distinct signaling events to eliminate the threat, including activation of IRF3 signaling, NF-κB signaling, and autophagy (1–5).

The combination of these signaling events triggered by STING can lead to drastically different cellular outcomes depending on the context. For example, while interferon production through activation of IRF3 has potent anti-viral and anti-cancer effects, it has been shown that interferon can actually increase the virulence of some bacterial pathogens, such as *Listeria monocytogenes* (6, 7). In addition, chronic elevated interferon production in the absence of a real threat can lead to debilitating autoimmune conditions, such as systemic lupus encephalitis, and neurodegeneration (8–10). Although STING is activated in many different contexts, it is unknown if its downstream signaling events can be modulated to ensure a context-appropriate response to a particular threat.

Here we identify PELI2, an E3 ubiquitin ligase, as a regulator of STING signaling outcomes. PELI2 selectively inhibits STING-mediated activation of IRF3 while enhancing activation of NF-κB, tuning the balance between these signaling outcomes. Mechanistically, we show that PELI2 is recruited to phosphothreonine residues on STING where it can target the kinase TBK1 for ubiquitination to reduce IRF3 phosphorylation and activation. While under basal conditions, PELI2 suppresses STING activation of interferons to prevent autoimmunity, we find that interferons downregulate PELI2, creating a positive feedback loop that allows STING to be fully active when a threat is present.

## Results

### PELI2 selectively inhibits the IRF3 axis of STING signaling, while enhancing the NF-κB axis

We previously performed genome-wide CRISPR screens to identify regulators of cGAMP-STING signaling in U937 human monocytic cells (11, 12). One of the highest ranked negative regulators from these screens is the E3-ubiquitin ligase PELI2. While PELI2 has not previously been suggested to be involved in STING signaling, studies have shown that PELI2 positively regulates other immune pathways, including TLR9 signaling and NLRP3 inflammasome activation (13, 14). To validate PELI2 as a negative regulator of STING signaling, we transduced U937 cells with a vector expressing Cas9 and a PELI2 sgRNA to create a PELI2 knockout cell line. Although STING signaling is multibranched and activates multiple distinct pathways, our CRISPR screens enriched selectively for regulators of the IRF3 axis of STING signaling. We thus first focused on determining PELI2’s effects on the STING-IRF3 signaling axis. Mechanistically, the sequence of events that lead to activation of the transcription factor IRF3 through STING signaling have been well characterized: first, cGAMP binding to STING leads to a conformational change that exposes STING’s C-terminal tail (CTT); second, the kinase TBK1 binds to a motif at the end of the exposed STING CTT and becomes active through auto-phosphorylation; third, active TBK1 phosphorylates the STING CTT on multiple serine and threonine residues, including Ser366; finally, IRF3 binds to phosphorylated Ser366 on the STING CTT, positioning IRF3 to be phosphorylated and activated by TBK1.

We evaluated the effect of PELI2 on each of these steps by measuring phosphorylation levels of TBK1, STING, and IRF3 following cGAMP treatment in WT or PELI2 knockout cell lines. Interestingly, cGAMP-treated PELI2 knockout cells had significantly more IRF3 phosphorylation than WT cells (**Figure S1a-b**); however, STING and TBK1 phosphorylation did not differ between the two cell lines (**Figure S1c-d**). To see if this effect could be rescued by expression of ectopic PELI2, we transduced a doxycycline-inducible, sgRNA-resistant PELI2-FLAG expression vector into U937 PELI2 sgRNA cells to create U937 tet-PELI2-FLAG cells.

Using these cells, we found that induction of PELI2 reduces IRF3 phosphorylation in response to cGAMP by ∽50% compared to the PELI2 knockout cells (**Figure 1a-b**). Furthermore, PELI2 also reduced levels of STING phosphorylation by ~25%, while having no effect on levels of TBK1 phosphorylation (**Figure 1c-d**). Together, these data indicate that PELI2 negatively regulates the IRF3 axis of STING signaling.

**Figure 1:**
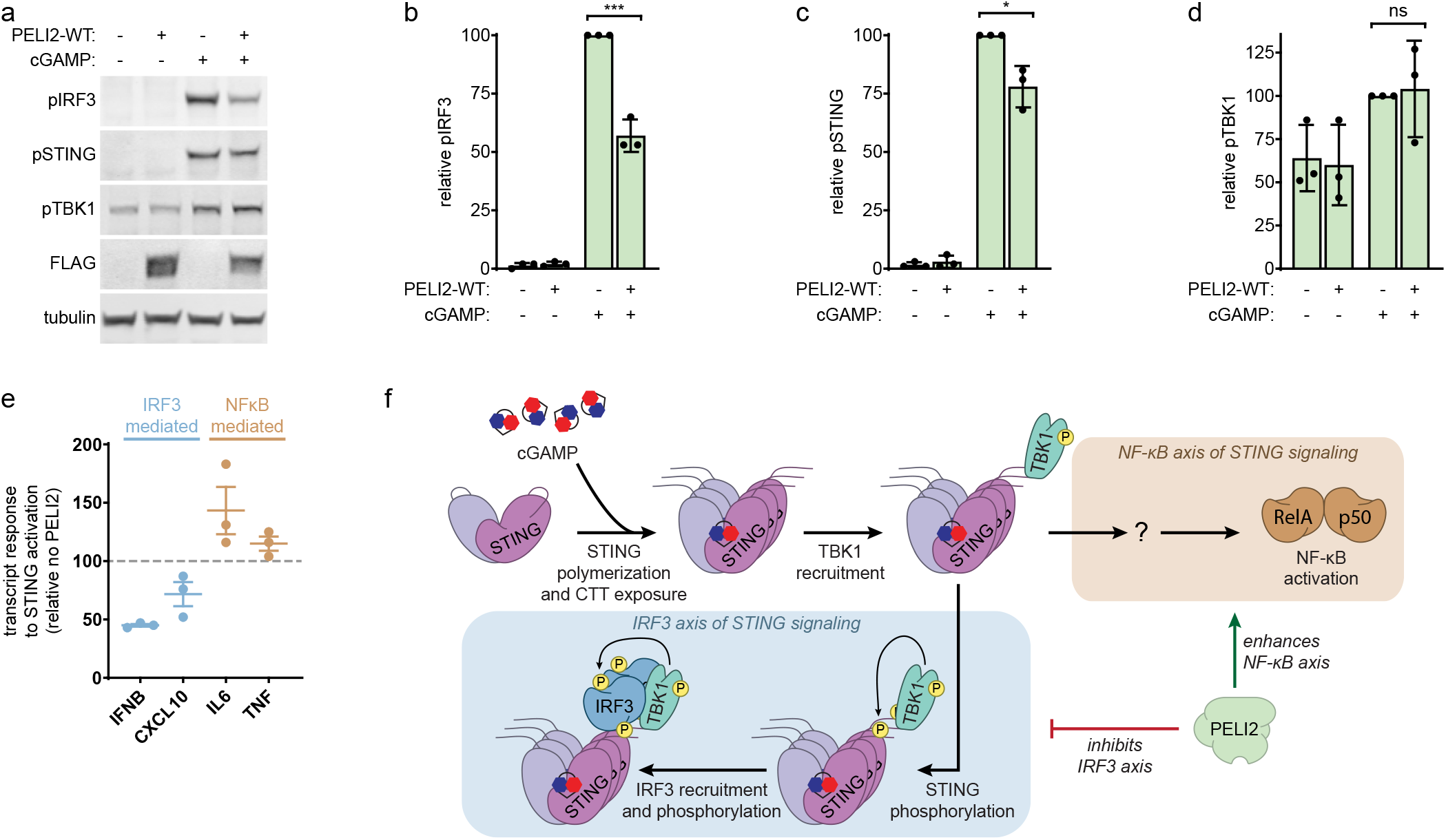
PELI2 selectively inhibits the IRF3 axis of STING signaling, while enhancing the NF-κB axis. a) U937 tet-PELI2-FLAG-WT cells were treated with 20 ng / mL doxycycline overnight to induce PELI2-FLAG expression before treatment with 50 μM cGAMP for 2 h. Cell lysates were then processed for Western blot analysis. Band intensity was quantified and normalized to tubulin for b) phosphorylated IRF3, c) phosphorylated STING, and d) phosphorylated TBK1. e) U937 tet-PELI2-FLAG-WT cells were induced with 20 ng / mL doxycycline overnight before treatment with 100 μM cGAMP for 3 h. Samples were then processed for RT-qPCR analysis of IFNB, CXCL10, IL6, and TNF transcript levels. Shown are transcript levels in PELI2 induced cells relative to uninduced cells. f) Model for PELI2’s regulation of STING signaling.

Since PELI2 and its family members have been previously implicated in indirect positive regulation of the NF-κB pathway through their effects on TLR signaling (13, 15), we next investigated whether PELI2 had a similar effect on NF-κB activation through the STING pathway. To do this, we treated U937 tet-PELI2-WT cells and measured the effect of ELI2 induction on the NF-κB-controlled transcripts IL6 and TNF by RT-qPCR following cGAMP stimulation. While PELI2 induction reduced levels of IRF3-controlled transcripts IFNB and CXCL10, it increased levels of NF-κB-regulated transcripts in response to cGAMP (**Figure 1e**). This suggests that PELI2 selectively inhibits the IRF3 axis of STING signaling, while enhancing the NF-κB axis (**Figure 1f**).

### PELI2 phosphothreonine binding is necessary for inhibition of the STING-IRF3 axis

Since the IRF3 axis of STING signaling has been mechanistically well characterized, while the NF-κB axis has not, we decided to focus our efforts on uncovering the molecular details of how PELI2 specifically inhibits IRF3 activation. PELI2 consists of two known functional domains: an N-terminal FHA domain that binds to phosphothreonine residues and a C-terminal RING domain that facilitates ubiquitination of target substrates. A mutation in the FHA domain, R106A, has previously been shown to prevent binding to phosphothreonine (**Figure 2a**) (16). To assess whether PELI2’s phosphothreonine binding activity is necessary for inhibition of the STING-IRF3 axis, we inserted a dox-inducible, sgRNA-resistant PELI2-FLAG-R106A expression vector into U937 PELI2 sgRNA cells. In contrast to expression of PELI2-WT, induction of PELI2-R106A did not inhibit IRF3 or STING phosphorylation in response to cGAMP (**Figure 2b-e)**. These results suggest that PELI2 binds to some unknown phosphothreonine residue as a requisite step in its inhibition of the STING-IRF3 axis.

**Figure 2:**
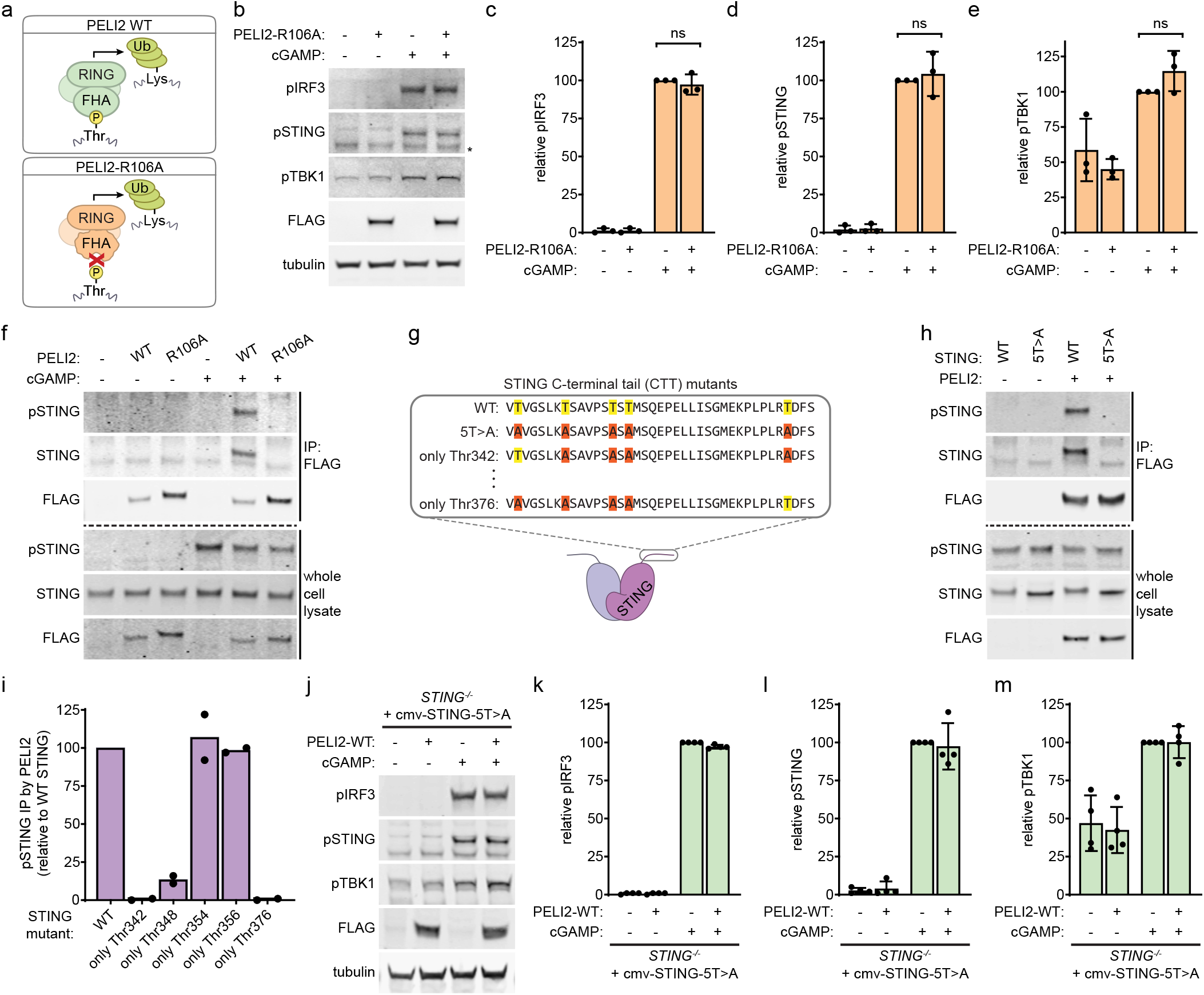
PELI2 binds to phosphorylated Thr354 and Thr356 on STING’s CTT to inhibit IRF3 activation. a) Schematic of domain composition and function of PELI2 WT and R106A. b) U937 tet-PELI2-FLAG-R106A cells were treated with 20 ng / mL doxycycline overnight to induce PELI2-FLAG-R106A expression before treatment with 50 μM cGAMP for 2 h. Cell lysates were then processed for Western blot analysis. Band intensity was quantified and normalized to tubulin for c) phosphorylated IRF3, d) phosphorylated STING, and e) phosphorylated TBK1. f) U937 tet-PELI2-FLAG-WT and tet-PELI2-FLAG-R106A cells were induced with 20 ng / mL doxycycline overnight before treatment with 50 μM cGAMP for 2 h. PELI2-FLAG was then immunoprecipitated from whole cell lysate with anti-FLAG beads. Western blot analysis was performed on immunoprecipitated FLAG and whole cell lysate. g) Schematic of CTT sequence of STING WT and STING CTT mutants. WT threonines are highlighted in yellow, whereas threonines mutated to alanine are highlighted in red. h) HEK293T cells were transfected with PELI2-FLAG-WT and either STING-WT or STING-5T>A. Then, 2 d post transfection, cells were treated with 100 μM cGAMP for 2 h before lysing cells for FLAG immunoprecipitation and subsequent Western blot analysis. i) HEK293T cells were transfected with PELI2-FLAG WT and either STING-WT or various STING CTT mutants. Then, 2 d post transfection, cells were treated with 100 μM cGAMP for 2 h before lysing cells for FLAG immunoprecipitation and subsequent Western blot analysis. Levels of immunoprecipitated phosphorylated STING mutants were quantified and normalized to levels of immunoprecipitated phosphorylated STING-WT. j) U937 STING^-/-^ + cmv-STING-5T>A + tet-PELI2-FLAG-WT cells were induced with 20 ng / mL doxycycline overnight before treatment for 2 h. Band intensity was quantified and normalized to tubulin for k) phosphorylated IRF3, l) phosphorylated STING, and m) phosphorylated TBK1.

### PELI2 binds to phosphothreonines on the C-terminal tail of STING to inhibit IRF3 activation

Given that IRF3 is activated on STING’s CTT, which contains multiple potential phosphothreonine sites, the STING CTT appeared to be a likely candidate for the binding site of PELI2. We first tested whether PELI2 interacts with full-length STING through immunoprecipitation assays of PELI2-FLAG from lysates of U937 tet-PELI2-FLAG cells. Pulldown of unphosphorylated STING by anti-FLAG beads was the same in both negative control and PELI2-FLAG expressing cells, indicating that PELI2 does not interact with unphosphorylated STING. In cells treated with cGAMP, STING phosphorylation can be detected both with an antibody that specifically recognizes phosphorylation of Ser366 on STING and with the appearance of a higher molecular weight phosphorylated STING band above the normal, unphosphorylated STING band. With both of these detection methods, we observed that phosphorylated STING is pulled down specifically by WT PELI2-FLAG, but not by R106A PELI2-FLAG or negative control cells (**Figure 2f**). Together, these data indicate that PELI2 binds to phosphorylated STING.

The STING CTT contains five threonine residues that, once phosphorylated, could potentially act as binding sites for PELI2 (**Figure 2g**). To determine if these threonines are necessary for PELI2’s interaction with STING, we created a STING mutant where all five threonines are mutated to alanine (STING-5T>A) and coexpressed it with PELI2-FLAG in HEK293T cells for co-IP analysis. While wild-type STING strongly co-precipitated with PELI2-FLAG, STING-5T>A did not. Importantly, phosphorylation of Ser366 was not diminished for STING-5T>A, demonstrating that the lack of PELI2 interaction observed for this mutant is not due to an overall defect in STING phosphorylation at other sites (**Figure 2h**). To determine which threonines alone are sufficient to enable PELI2 binding to phosphorylated STING, we created an additional five STING mutants in which all but one of the threonines in the CTT are mutated to alanine. We found that STING mutants containing either only Thr354 or only Thr356 were both sufficient to completely rescue pulldown by PELI2-FLAG (**Figure 2i, Figure S2a)**. Together, these results demonstrate that PELI2 can bind either phosphorylated Thr354, Thr356, or both on the CTT of activated STING.

To validate that PELI2 binding to STING is necessary for its inhibition of STING-IRF3 signaling, we created a U937 cell line with endogenous STING knocked out and replaced with STING-5T>A driven by the CMV promoter, then transduced our tet-PELI2-FLAG-WT expression vector into these cells. Activation of STING-IRF3 signaling by these STING-5T>A cells appears to be unimpaired, as these cells still phosphorylate TBK1, STING, and IRF3 in response to cGAMP stimulation. In contrast to cells with endogenous STING, however, PELI2 expression had no effect on the STING-IRF3 signaling axis in these STING-5T>A cells (**Figure 2j-m**), showing that PELI2 binding to STING is necessary for its inhibition of the STING-IRF3 axis.

### The RING domain of PELI2 is required for its inhibition of STING-IRF3 signaling

Given the close proximity of the PELI2 binding site on the STING CTT to the IRF3 binding site, it is possible that PELI2 inhibits IRF3 activation by sterically blocking IRF3 from binding to STING. Alternatively, we considered that PELI2 binding to STING could bring PELI2 to close proximity with potential ubiquitination substrates like TBK1 or IRF3 – whose ubiquitination and degradation would be expected to inhibit STING-IRF3 signaling. To distinguish between these two possibilities, we took advantage of the previously described PELI2-C397/400A mutant (hereafter referred to as PELI2-CC>AA), which has a defective E3 ubiquitin ligase RING domain (**Figure 3a**) (17). After inserting our sgRNA-resistant tet-PELI2-FLAG-CC>AA expression vector into U937 PELI2 sgRNA cells, we found that PELI2-CC>AA retained the ability to pull down phosphorylated STING, indicating that it still has a functional FHA domain (**Figure S3a**). PELI2-CC>AA induction, however, did not inhibit phosphorylation of IRF3, STING, or TBK1 in response to cGAMP, indicating that a functional RING domain is required for inhibition of the STING-IRF3 axis (**Figure 3b-e**). Supporting this notion, induction of a truncated PELI2 containing only the FHA domain was also unable to inhibit phosphorylation of IRF3, STING, or TBK1 (**Figure S3b-e**), indicating that PELI2 binding to STING alone is not sufficient to inhibit STING-IRF3 signaling.

**Figure 3:**
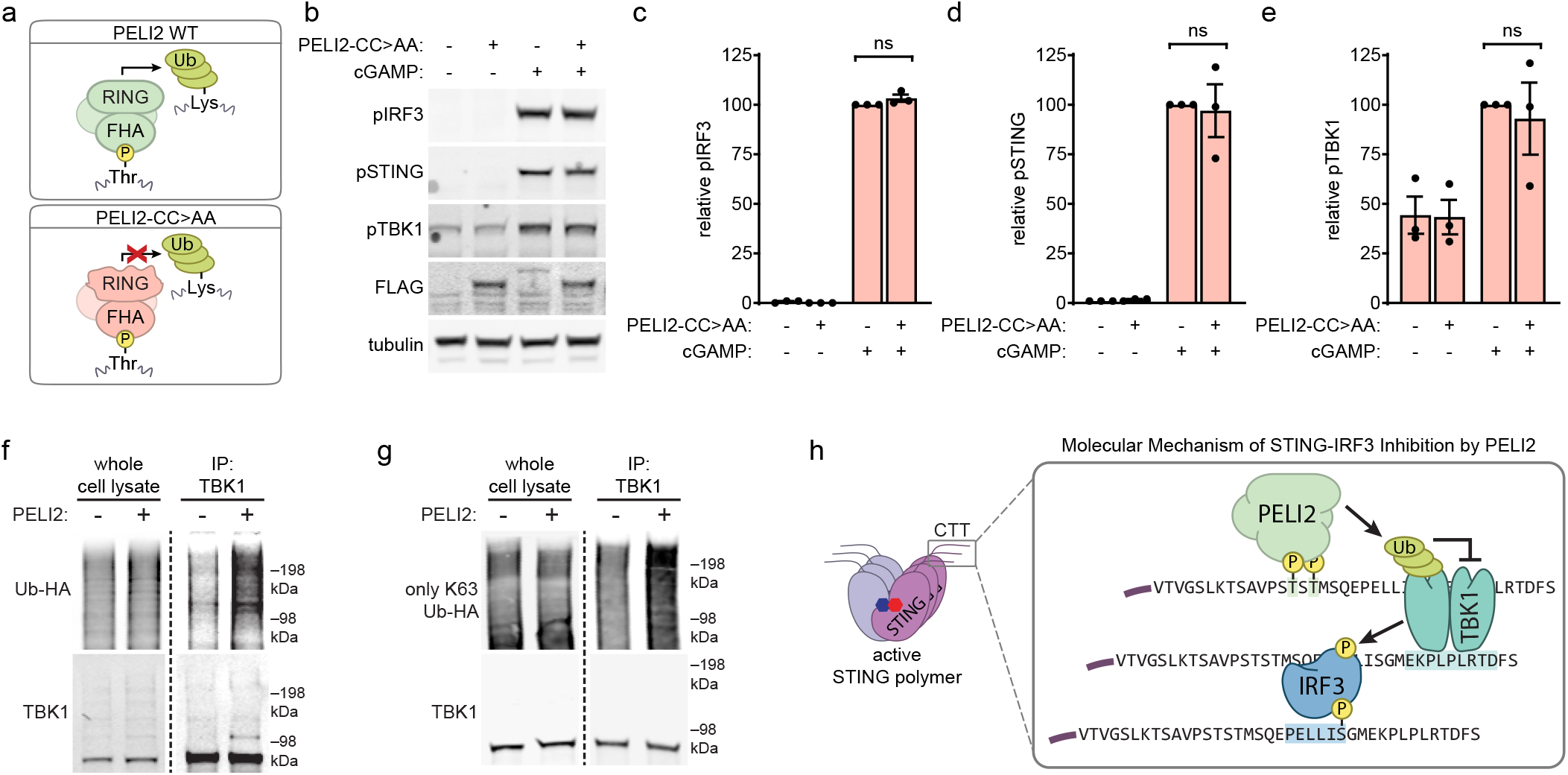
The RING domain of PELI2 promotes ubiquitination of TBK1 and is necessary for STING-IRF3 inhibition. a) Schematic of domain composition and function of PELI2 WT and CC>AA. b) U937 tet-PELI2-FLAG-CC>AA cells were treated with 20 ng / mL doxycycline overnight to induce PELI2-FLAG-CC>AA expression before treatment with 50 μM cGAMP for 2 h. Cell lysates were then processed for Western blot analysis. Band intensity was quantified and normalized to tubulin for c) phosphorylated IRF3, d) phosphorylated STING, and e) phosphorylated TBK1. f) HEK293T cells were transfected with or without PELI2-FLAG-WT and with HA tagged ubiquitin (Ub-HA). Then, 2 d post transfection, cells were treated with 100 μM cGAMP for 2 h before lysing cells for TBK1 immunoprecipitation. Western blot analysis was performed on immunoprecipitated TBK1 and whole cell lysate samples. g) Similar to f), except cells were transfected with an Ub-HA mutant where all lysines except K63 are mutated to arginine (only K63 Ub-HA). h) Model for the molecular mechanism of STING-IRF3 inhibition by PELI2.

Since the E3 ubiquitin ligase activity of PELI2 is required for its inhibition of STING-IRF3 signaling, we next investigated what target(s) PELI2 could be ubiquitinating to exert these effects. We used HEK293T cells expressing HA-tagged ubiquitin together with a transfected STING-FLAG construct to measure ubiquitination levels of endogenous IRF3, endogenous TBK1, or heterologous STING-FLAG. While overexpression of PELI2 had no effect on ubiquitination levels of IRF3 or STING (**Figure S3f-g**), we found a notably increase in ubiquitination of endogenous TBK1 upon PELI2 overexpression (**Figure 3f**).

Since PELI family members have been suggested to be capable of forming both K48 and K63 linked ubiquitin chains, we sought to determine which type of ubiquitin chain PELI2 adds to TBK1. To do this, we took advantage of ubiquitin mutants capable of forming either only K48- or only K63-linked chains. While PELI2 had no effect on TBK1 ubiquitination by Ub-HA capable of forming only K48 linkages (**Figure S3h**), PELI2 still increased TBK1 ubiquitination by Ub-HA capable of forming only K63 linkages (**Figure 3g**). Together, these results suggest the following model for how PELI2 inhibits STING-IRF3 signaling: after TBK1 phosphorylates the STING CTT, it creates a binding site for PELI2 at phospho-Thr354/356; PELI2 bound to STING CTT is brought in close proximity to TBK1, leading to its K63-linked ubiquitination; K63-linked ubiquitination of TBK1 results in its decreased activity toward phosphorylating STING and IRF3 (**Figure 3h**). Since K63-linked ubiquitination has previously been implicated in activating NF-κB signaling (18), this could also provide a possible explanation for how PELI2 increases NF-κB signaling.

### PELI2 prevents basal STING activation from leading to elevated interferon signaling

While the NF-κB axis of STING signaling is evolutionarily ancient and is observed in animals as distantly related from humans as sea anemones, the STING-IRF3 axis of signaling evolved more recently within vertebrates (19). Although the STING-IRF3 axis is a powerful tool for anti-viral and anti-cancer immunity, prolonged activation of STING-IRF3 can lead to interferonopathy and serious autoimmune conditions, including SAVI and systemic lupus encephalitis (SLE) (8–10, 20). SLE is characterized by chronic elevated interferon signaling resulting in upregulation of interferon stimulated genes (ISGs). Since our previous experiments demonstrated that PELI2 suppresses STING-IRF3 signaling following short, high dose cGAMP stimulation, we next sought to determine whether PELI2 also suppresses chronic low levels of STING signaling that can lead to autoimmune conditions. U937 cells are cancerous and likely have some basal STING activation as a result, so we first assessed transcript levels of a panel of ISGs in U937 cells expressing Cas9 and either scramble sgRNA, PELI2 sgRNA, or STING sgRNA under basal conditions. Remarkably, we found that ISG expression levels were consistently higher in PELI2 sgRNA cells compared to scramble sgRNA cells. We found that all but one of the measured ISGs were downregulated in STING^-/-^ cells, supporting the idea of basal STING activation in these cells that is suppressed by PELI2 (**Figure 4a**). To validate that PELI2’s suppression of ISGs is occurring through STING, we used U937 tet-PELI2 cells with WT or 5T>A STING. Strikingly, PELI2 reduced ISG levels by ~90% in WT STING cells following a prolonged, low dose cGAMP treatment; in contrast, PELI2 had little to no effect on ISG levels in 5T>A STING cells under the same conditions (**Figure 4b**). Together, these results indicate that PELI2 is able to dampen IRF3 signaling in conditions of chronic STING activation, in addition to inhibition of acute STING activation.

**Figure 4:**
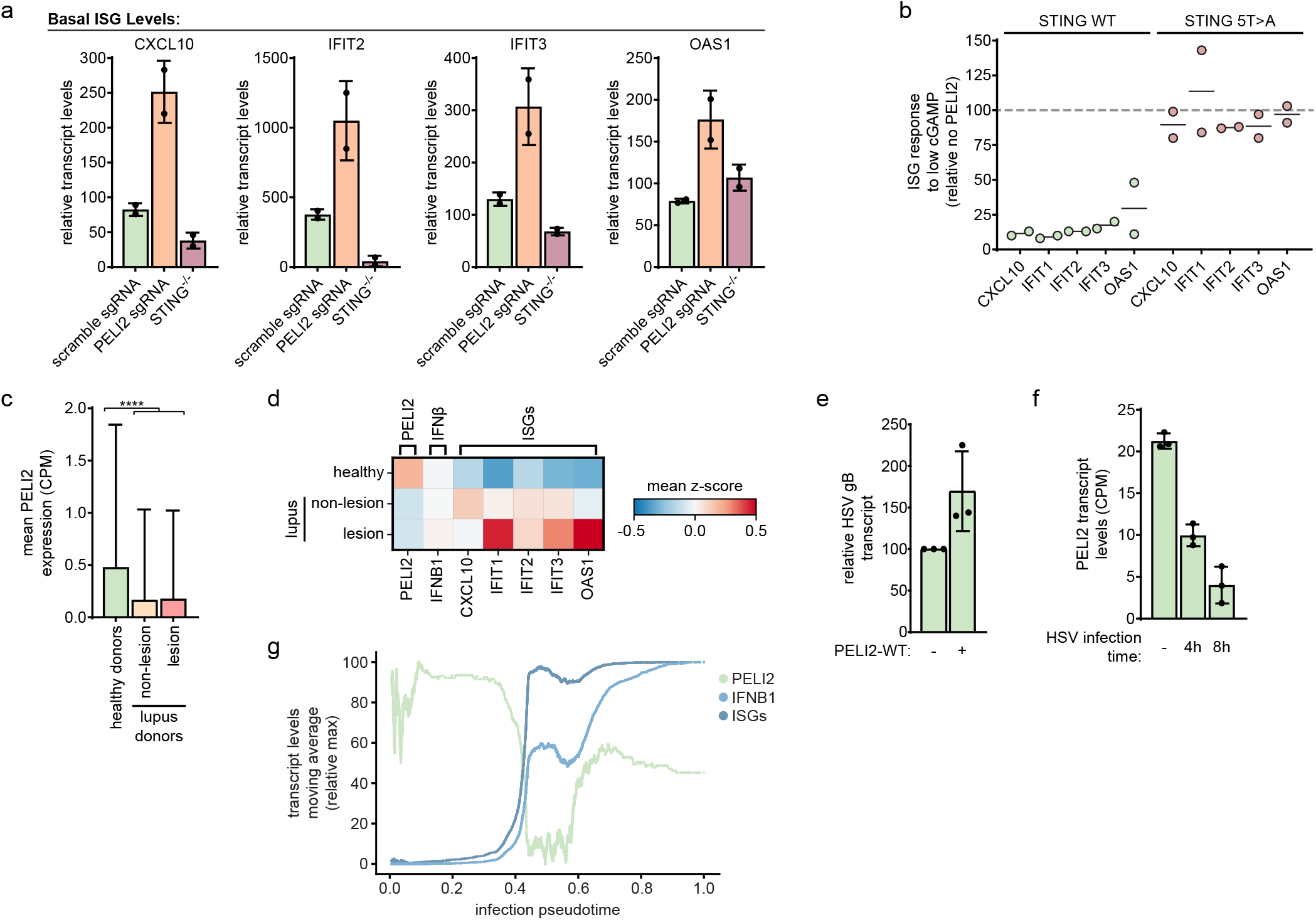
PELI2 suppresses interferon signaling from basal STING activation and is itself downregulated by interferon. a) Basal levels of ISGs in U937 scramble sgRNA, PELI2 sgRNA, or STING^-/-^ cells were analyzed by RT-qPCR. b) U937 tet-PELI2-FLAG-WT cells with endogenous STING and U937 STING^-/-^ + cmv-STING-5T>A + tet-PELI2-FLAG-WT cells were induced with 20 ng / mL doxycycline overnight before treatment with low dose cGAMP (10 μM) for 23 h. Levels of ISGs were analyzed by RT-qPCR and are shown relative to uninduced cells. c) Single cell RNA-seq data showing mean expression of PELI2 in keratinocytes from healthy donors or lupus patients. Skin samples were collected from both non-lesional and lesional sites for lupus patients (raw data from GSE186476). d) Heatmap showing relative expression of PELI2, IFNB1, and ISGs in keratinocytes from healthy donors and lupus patients. e) MDA-MB-231 tet-PELI2-FLAG-WT cells were induced with 20 ng / mL doxycycline overnight before infection with HSV at 1 MOI. Then, 6 hours post infection, cells were lysed and HSV gB transcript levels were measured by RT-qPCR. f) RNA-seq quantification of PELI2 transcript levels in HSV infected cells (GSE124118). HFF-1 cells were mock infected or infected with HSV at 0.5 MOI for 4 or 8 h. g) Pseudotime analysis of PELI2, IFNB1, and ISG transcript levels in plasmacytoid dendritic cells following infection with influenza virus.

Having shown that PELI2 suppresses interferon signaling through STING, we wondered if decreased expression of PELI2 could increase risk of autoimmunity. The majority of patients with SLE develop skin lesions driven by elevated interferon signaling following UV exposure, which can activate STING (21, 22). Given that keratinocytes have been suggested to be the major initial source of elevated interferons in these lesions (23), we investigated PELI2 levels in keratinocytes from skin samples of healthy donors and lupus patients from a previously published single-cell RNA sequencing dataset (24). After cell type clustering (**Figure S4a**), we found that PELI2 expression was over 50% lower in the keratinocytes of lupus patients compared to healthy donors (**Figure 4c**). Interestingly, these lower levels of PELI2 were observed in non-lesional samples from lupus patients as well as lesional samples. Furthermore, there was a reverse correlation between PELI2 expression and expression of IFNβ and ISGs in these keratinocytes (**Figure 4d, S4b**). Together, these results suggest that reduced PELI2 levels in lupus patients could contribute to their hypersensitivity towards UV light.

### PELI2 dynamically regulates STING-IRF3 during viral infection

The anti-viral properties of STING signaling have been well characterized, particularly with regard to the dsDNA virus Herpes Simplex Virus 1 (HSV). Since PELI2 inhibits STING-IRF3 signaling, it seemed likely that PELI2 would consequently also inhibit the anti-viral response to HSV. To test this, we infected tet-PELI2-WT cells with HSV and assessed transcript levels of the HSV late gene gB to measure viral replication. We found that PELI2 overexpression increased HSV gB transcript levels by over 50% (**Figure 4e**), indicating that PELI2 hinders the anti-viral response to HSV.

From our data so far, we can conclude that PELI2 plays an important role in the regulation of STING-IRF3 signaling in basal conditions and viral infection. While inhibiting ISG activation in the context of basal STING activation would be protective against autoimmunity, it is important that STING can be robustly activated in response to acute infection. We considered the possibility that PELI2 could be regulated to release the brake on STING-IRF3 signaling in certain conditions. To assess this, we analyzed published bulk RNA-seq data (GEO: GSE124118) of HFF-1 cells and found that PELI2 transcript levels drastically decrease following HSV infection (**Figure 4f**). Similarly, analysis of a separate dataset (GEO: GSE67983) shows that PELI2 transcript levels decrease in primary human monocytes following infection with the intracellular bacteria *Listeria monocytogenes* (**Figure S4c**). These associations support the notion that PELI2 could be downregulated in response to infection to allow for maximal STING activation.

To obtain a higher resolution depiction of the dynamics of PELI2 and IFNβ transcript levels following viral infection, we performed diffusion pseudotime analysis on published data from primary plasmacytoid dendritic cells infected with influenza virus (25). Viral treatment time correlated well with pseudotime, validating that pseudotime analysis is capturing the transcriptional response to viral infection over time (**Figure S4d**). Consistent with the bulk RNA-seq analysis above, we found that PELI2 levels rapidly decrease by ~80% following viral infection. This decrease coincides with an initial increase in IFNB transcript expression, which then starts to plateau. Interestingly, after PELI2 levels reach a minimum, IFNB transcript levels start to increase even further, supporting a model where downregulation of PELI2 initiates a positive feedback loop to potentiate IFNβ signaling. Furthermore, PELI2 expression appears to be restored late in viral infection, resetting a basal-like state (**Figure 4g**).

Given that HSV infection results in cGAMP production (26), we reasoned that the observed downregulation of PELI2 transcript may actually lie downstream of the STING pathway itself. Indeed, we found that PELI2 transcript levels were downregulated in WT U937 cells following treatment with a high-dose of cGAMP. Interestingly, this effect was not observed in U937 IRF3 sgRNA cells, indicating that PELI2 downregulation depends on activation of the STING-IRF3 axis. Furthermore, PELI2 downregulation was only observed after 6 h of cGAMP stimulation, with no effect at the earlier 3 h timepoint (**Figure S4e**). Supporting the idea that a particular threshold of STING signaling must be reached to initiate PELI2 downregulation, we did not observe PELI2 downregulation in U937 cells where STING was weakly stimulated with a low concentration of cGAMP for a prolonged time (**Figure S4f**). Since induction of transcripts directly controlled by IRF3, such as IFNB, can be observed as early as 3 h after cGAMP treatment, this delayed response suggests that PELI2 may be an indirect target or further downstream in the IRF3 regulon, rather than a primary target like IFNβ. In fact, we found that IFNβ treatment alone was sufficient to decrease PELI2 levels (**Figure S4g**), suggesting that PELI2 is a secondary target downstream of the STING-IRF3-IFNβ axis.

Together, our data suggest that under basal conditions, PELI2 is important for suppressing low levels of STING activation. However, once STING activation surpasses a certain threshold, enough IFNβ is produced to downregulate PELI2, priming cells for possible threats by increasing the sensitivity of the STING pathway.

## Discussion

Over the past decade, the STING pathway has emerged as a central pathway in immunity implicated in a plethora of diseases and conditions, including viral infection, cancer, neurodegeneration, heart attack, and autoimmunity (10, 27–31). The STING pathway must be properly regulated to ensure a balance between tolerating weak stimuli of pathway activation that may occur basally, while decisively responding to strong stimuli of pathway activation that occur during cellular threats such as viral infection. Furthermore, given the wide array of conditions in which STING activation can occur, the downstream effects of the pathway, such as IRF3 and NF-κB signaling, must be differentially regulated to ensure a context appropriate response. However, such regulatory mechanisms have remained elusive. Here, we have uncovered PELI2 as a key regulator of balancing both the strength and effects of STING activation.

Mechanistically, PELI2 inhibits STING-IRF3 by binding to a phosphorylated site on STING’s CTT and subsequently ubiquitinating TBK1, thereby constraining its activation of IRF3.Since NF-κB signaling can be activated by K63 linked ubiquitin chains, it’s possible that PELI2’s ubiquitination of TBK1 also promotes NF-κB activation. While PELI2 inhibits interferon signaling through STING, PELI2 is itself downregulated through interferon signaling. Our data supports a model in which PELI2 suppresses STING-IRF3 activation under basal conditions, but once a threshold of interferon signaling has been reached, PELI2 is downregulated to fully unleash the STING pathway.

While many different genetic dispositions and environmental factors can lead to autoimmune conditions such as SLE, at its essence SLE is a disease driven by dysregulated, overactive interferon signaling. We have shown that PELI2 is downregulated in skin keratinocytes of SLE patients, which could contribute to SLE patients’ STING-dependent sensitivity to UV light. Given PELI2’s central role in modulating STING signaling and basal interferon production, we posit that variations in PELI2 function and regulation across the population could contribute to the risk of SLE and other interferonopathies.

In mammals, the CTT of STING plays an essential role in the activation of interferon signaling by recruiting TBK1 and IRF3 through specific motifs. Interestingly, in STING homologs of other vertebrates, such as zebrafish, the CTT contains alternative motifs that recruit other effectors instead of TBK1 and IRF3. Because of this, zebrafish STING primarily activates NF-κB signaling instead of interferon signaling (32). This indicates that the composition of STING’s CTT has a strong influence on the effects of STING signaling, and that different vertebrate species have evolved distinct CTT compositions to best suit their needs. With our identification of the PELI2 recruitment motif on STING’s CTT, we have identified another mechanism through which the CTT can be decorated to modulate STING signaling.

While most mammals possess at least one of the threonines important for PELI2 recruitment in human STING, some clades have no predicted PELI2 recruitment site (e.g. the Muridae family, containing mice). Furthermore, only a few clades contain a redundant threonine recruitment site like that observed in human STING (e.g. the Simiiformes infraorder of primates) (**Figure S5**). One possible explanation for these differences is that certain species, such as mice, may have lost the PELI2 recruitment site to evolve a STING pathway that is more sensitive to pathogens; alternatively, other species, such as humans, may have gained an additional PELI2 recruitment site to evolve a STING pathway that is more selective to stimulation. In fact, the human STING pathway has been shown to be less reactive/more selective than that of mice at two nodes: 1) mouse STING binds bacterial cyclic-di-nucleotides, cyclic-di-GMP and cyclic-di-AMP, with much higher affinity than human STING(33, 34); and 2) mouse cGAS is more sensitive to stimulation by dsDNA, whereas human cGAS is more selective for longer molecules of dsDNA and exhibits slower cGAMP synthesis kinetics (35). Here, our report of PELI2 regulation of human but not mouse STING adds another mechanism of evolutionary divergence that renders the human cGAS-STING pathway less reactive to pathogens, and also more tolerant to self-DNA.

PELI2 has previously been demonstrated to positively regulate multiple immune pathways, including NLRP3 inflammasome activation, TLR9 signaling, and IL-1R signaling (13, 14, 17). With our study, we provide the first evidence that PELI2 can also negatively regulate immune signaling. The role of PELI2 in differentially regulating immune pathways suggests it may influence the overall immune response to threats. Since threats can such as bacterial infection activate multiple distinct immune pathways, the presence of PELI2 may allow for the preferential activation of certain pathways (e.g. TLR signaling) over others (e.g. STING signaling). Given that different immune pathways elicit distinct immune responses, balancing activation of these different pathways through PELI2 could allow cells to ensure a context appropriate response to threats. Together, since PELI2 appears to regulate multiple immune pathways that have distinct signaling consequences, PELI2 levels in different cell types may not only determine the threshold and strength of a cell’s immune response to threats, but also the type of immune response.

Taken together, our study establishes PELI2 as a key regulator of STING signaling, not only biasing toward NF-κB pathway outputs and away from the IRF3 pathway, but also dynamically tuning the STING activation threshold to appropriately respond to a wide range of conditions including basal activity, chronic low-grade stimulation, acute viral infection, and recovery after viral infection. PELI2 is therefore a critical regulatory mechanism for human STING, and future studies will further illuminate its contributions to autoimmunity, viral infection, and many other contexts in which STING signaling is implicated.

## Acknowledgements

We would like to thank all Li lab members and B. Xu Hua Fu for their insightful comments that helped guide this study. This study was funded by NIH Grant 5T32GM007276 and the Arc Institute. C.R. was supported by the Stanford Graduate Fellowship.

## Figure Captions

**Supplemental Figure 1: PELI2 selectively inhibits the IRF3 axis of STING signaling, while enhancing the NF-**κ**B axis**.

a) U937 WT and PELI2 sgRNA cells were treated with 50 μM cGAMP for 2 h. Cell lysates were then processed for Western blot analysis. Band intensity was quantified and normalized to tubulin for b) phosphorylated IRF3, c) phosphorylated STING, and d) phosphorylated TBK1.

**Supplemental Figure 2: PELI2 binds to phosphorylated Thr354 and Thr356 on STING’s CTT to inhibit IRF3 activation**.

a) HEK293T cells were transfected with PELI2-FLAG WT and either STING-WT or various STING CTT mutants. Then, 2 d post transfection, cells were treated with 100 μM cGAMP for 2 h before lysing cells for FLAG immunoprecipitation and subsequent Western blot analysis.

**Supplemental Figure 3: The RING domain of PELI2 promotes ubiquitination of TBK1 and is necessary for STING-IRF3 inhibition**.

a) U937 tet-PELI2-FLAG-CC>AA cells were induced with 20 ng / mL doxycycline overnight before treatment with 50 μM cGAMP for 2 h. PELI2-FLAG was then immunoprecipitated from whole cell lysate with anti-FLAG beads. Western blot analysis was performed on immunoprecipitated FLAG and whole cell lysate.

b) U937 tet-PELI2-FLAG-FHA-only cells were treated with 20 ng / mL doxycycline overnight before treatment with 50 μM cGAMP for 2 h. Cell lysates were then processed for Western blot analysis. Band intensity was quantified and normalized to tubulin for c) phosphorylated IRF3, d) phosphorylated STING, and e) phosphorylated TBK1.

f) HEK293T cells were transfected with or without PELI2-FLAG-WT and with HA tagged ubiquitin (Ub-HA). Then, 2 d post transfection, cells were treated with 100 μM cGAMP for 2 h before lysing cells for IRF3 immunoprecipitation. Western blot analysis was performed on immunoprecipitated IRF3 and whole cell lysate samples.

g) HEK293T cells were transfected with Ub-HA, with or without STING-FLAG, and with or without PELI2-FLAG. Then, 2 d post transfection, cells were treated with or without 100 μM cGAMP for 2 h before lysing cells for FLAG immunoprecipitation. Western blot analysis was performed on immunoprecipitated FLAG and whole cell lysate samples.

h) HEK293T cells were transfected with or without PELI2-FLAG-WT and with a Ub-HA mutant where all lysines except K48 are mutated to arginine (only K48 Ub-HA). Then, 2 d post transfection, cells were treated with 100 μM cGAMP for 2 h before lysing cells for TBK1 immunoprecipitation. Western blot analysis was performed on immunoprecipitated TBK1 and whole cell lysate samples.

**Supplemental Figure 4: PELI2 suppresses interferon signaling from basal STING activation and is itself downregulated by interferon**.

a) Cell type clustering of single cell RNA-seq data of skin samples from healthy donors and lupus patients (raw data from GSE186476).

b) Expression of IFNB1 in keratinocytes of healthy donors and lupus patients.

c) Relative transcript levels of PELI2 in PBMCs infected with *Listeria monocytogenes* (5 MOI) for the indicated times (raw data from GSE67983).

d) Distribution of viral infection treatments times across pseudotime.

e) U937 WT and IRF3 sgRNA cells were treated with 100 μM cGAMP for 3 h or 6 h before analysis by RT-qPCR.

f) U937 WT cells were treated with 50 ng / mL IFNβ for 3 h before RT-qPCR analysis.

g) U937 WT cells were treated with a low dose of cGAMP (10 μM) overnight before RT-qPCR analysis of PELI2 transcript levels.

**Supplemental Figure 5: Phylogenetic tree of STING CTT evolution in mammals**

## Methods

### Cell Culture

HEK293T and MDA-MB-231 cells were maintained in DMEM supplemented with 10% FBS and 1% penicillin-streptomycin. U937 cells were maintained in RPMI supplemented with 10% heat-inactivated FBS and 1% penicillin-streptomycin. All cells were maintained in a 5% CO_2_ incubator at 37 °C.

### Creation of cell lines

pLentiCRISPR v2 was used as the 3rd-generation lentiviral backbone for all knockout lines. Guide sequences targeting PELI2, STING, or IRF3 were cloned into this lentiviral backbone using previously described protocols (36, 37).

pLVX-TetOne-FLAG was used as the lentiviral backbone for all inducible lines. PELI2-WT, PELI2-R106A, PELI2-2C>2A, and PELI2 FHA were cloned into this lentiviral backbone as previously described (12, 38).

pLenti-CMV-GFP-puro was used as the lentiviral backbone for creating stably expressing mutant STING lines. STING-5T>A was cloned into the BamHI-SalI site through Gibson assembly.

To produce lentivirus, HEK293T cells were transfected with 750 ng lentiviral plasmid, 750 ng pSPAX2, and 500 ng pHDM-G with FuGENE 6 transfection reagent. Viral media was exchanged after 24 h and harvested after 48 h. This viral media was then passed through a 0.45 μm filter before infection of cells. To infect U937 cells, 5 × 10^4^ cells were added to 1 mL viral media with 8 μg/mL polybrene. These cells were spun in a 24 well plate at 1000 × g for 1 h then resuspended in fresh media. To infect MDA-MB-231 cells, 5 × 10^4^ cells were split into a 12 well plate 24 h before infection. Then, media was removed from cells and replaced with 1 mL viral media. After a 24 h incubation, viral media was removed from cells and replaced with fresh media.

### cGAMP stimulation experiments for Western blot analysis

For experiments involving inducible U937 cell lines, cells were either left uninduced or induced with 20 ng / mL doxycycline overnight at a density of 5 × 10^5^ cells / mL before treatment. For all treatments, cells were pelleted and treated with the indicated concentration of cGAMP for 2 h at a density of 5 × 10^5^ cells / mL in 1.4 mL media. After treatment, cells were pelleted and lysed with 70 μL Laemmli Sample Buffer. Lysates were then sonicated and boiled for 10 min before running on a acrylamide gel for Western blot analysis.

### cGAMP stimulation experiments for RT-qPCR analysis

Cells were either left uninduced or induced with 20 ng / mL doxycycline overnight at a density of 5 × 10^5^ cells / mL before treatment. Cells were then pelleted and treated with the indicated concentration of cGAMP in the presence of 10 μM STF-1084 for the indicated amount of time at a density of 5 × 10^5^ cells / mL in 2 mL media. After treatment, cells were pelleted, wash with 1 mL PBS, and lysed with 350 μL TRIzol. RNA was purified from TRIzol using Direct-zol RNA miniprep kits (Zymo Research).

### Immunoprecipitation experiments

For immunoprecipitation experiments in U937 cells, cells were either left uninduced or induced with 20 ng / mL doxycycline at a density of 5 × 10^5^ cells / mL. Cells were then pelleted and treated with the indicated concentration of cGAMP for 2 h at a density of 1.5 × 10^6^ cells / mL in 1.4 mL media. Cells were then pelleted and lysed in 400 μL lysis buffer (Cell Signaling Technology) supplemented with protease and phosphatase inhibitors (Roche). Lysate was then clarified by centrifugation at 12,000 × g for 15 min.

For immunoprecipitation experiments in HEK293T cells, 1.5 × 10^5^ cells were split into a 6 well plate 24 h before transfection. For experiments investigating PELI2-FLAG pulldown of different STING mutants, cells were transfected with 20 ng pCMV-STING mutant, 2000 ng pcDNA3.1, and either 500 ng pcDNA3.1-PELI2-WT-FLAG or 500 ng pcDNA3.1 as a negative control. For experiments investigating ubiquitination of different proteins by PELI2, cells were transfected with 1000 ng pRK5-Ub-HA, 1800 pcDNA3.1, and either 200 ng pcDNA3.1-PELI2-WT-FLAG or 200 ng pcDNA3.1 as a negative control. For all HEK293T experiments, transfected cells were split into 10 cm dishes 24 h after transfection. Then, 48 h after transfection, cells were treated with 100 uM cGAMP for 2 h, washed with PBS, then lysed in 400 μL lysis buffer (Cell Signaling Technology) supplemented with protease and phosphatase inhibitors (Roche). Lysate was then clarified by centrifugation at 12,000 × g for 15 min.

For immunoprecipitation of PELI2-FLAG, 30 μL of anti-FLAG magnetic agarose (Pierce) was equilibrated 3× with lysis buffer. Then, 300 μL of clarified lysate was added to equilibrated agarose, and incubated for 20 min rotating at room temperature. Agarose was then washed 5× with 500 μL lysis buffer, transferred to a new tube, and eluted by boiling 2× in 35 μL Laemmli Sample Buffer for 5 min.

For immunoprecipiration of TBK1 and IRF3, 6 μL of either anti-TBK1 or anti-IRF3 was added to 294 μL clarified lysate and incubated overnight rotating at 4 °C. Then, 30 μL of protein A magnetic agarose (Cell Signaling Technology) was added incubated for an additional 4 h rotating at 4 °C. Agarose was then washed 5x with 500 μL lysis buffer, transferred to a new tube, and eluted by boiling 2x in 35 μL Laemmli Sample Buffer for 5 min.

### HSV infection

2 × 10^5^ MDA-MB-231 PELI2 sgRNA1 + tet-PELI2-FLAG WT cells were split into 12 well plate and either left uninduced or induced with 20 ng / mL doxycycline overnight. Then, cells were infected in 200 uL serum free DMEM with HSV-1 at 1 MOI. Plate was rocked every 15 min to ensure adequate coverage of the cells. After 1 h, virus was removed from the cells and replaced with 1 mL fresh, complete media. After an additional 6 h incubation, media was removed, and cells were lysed with 350 μL TRIzol. RNA was purified from TRIzol using Direct-zol RNA miniprep kits (Zymo Research).

### Analysis of bulk expression data

RNA-seq data for effects of HSV infection on transcript expression in HFF-1 cells were obtained from GEO, accession GSE124118. Counts of PELI2 transcript levels were normalized to counts per million (CPM) per sample.

Microarray data for effects of Listeria monocytogenes infection on transcript expression of primary monocytes was obtained from GEO, accession GSE67983. Quantile normalized values of probe ILMN_1780132 (which hybridizes to PELI2) were used to determine PELI2 expression values.

### Analysis of single cell RNA-seq data

Single cell RNA-seq data of skin biopsies from healthy donors and lupus patients were obtained from GEO, accession GSE186476 (24), and processed with scanpy version 1.9. Single cells with less than 500 genes detected or greater than 25% mitochondrial genes were filtered out prior to analysis. UMI counts were normalized to counts per million and log + 1 transformed. UMAPs of cells were created by running the following scanpy methods sequentially using the default parameters: pp.highly_variable_genes(), tl.pca(), pp.neighbors(), then tl.umap(). Leiden clustering was then performed on UMAPs by running tl.leiden() with resolution=0.1. Cell type annotations of Leiden clusters were determined by obtaining initial annotations with the CellO package (39), then manually updating these annotations to align with cell types known to be present in the skin. Cells annotated as keratinocytes were then selected to compare gene expression between patient conditions.

Single cell RNA-seq data of donor isolated plasmacytoid dendritic cells infected with influenza virus were obtained from GEO, accession GSE189120 (25), and processed with scanpy version 1.9. Single cells with less than 1000 genes, more than 6000 genes, or more than 20% mitochondrial genes were filtered out prior to analysis. UMI counts were normalized to counts per million and log + 1 transformed. UMAPs of cells were created by running the following scanpy methods sequentially using the default parameters: pp.highly_variable_genes(), tl.pca(), pp.neighbors(), then tl.umap(). Leiden clustering was then performed on UMAPs by running tl.leiden() with resolution=0.5. While the majority of cells clustered close together in UMAP space, a minority of cells appeared in a separate, distal cluster (cluster 5) and were filtered out prior to pseudotime analysis. Cells were assigned an ISG score using tl.score_genes() using the following list of 10 ISGs: IFIT1, IFIT2, IFIT3, CXCL10, OAS1, ISG15, ISG20, IRF7, IFI44L, and IFIH1. The cell with the lowest ISG score was designated as the root cell for pseudotime analysis. Diffusion pseudotime analysis was performed by sequentially running tl.diffmap() then tl.dpt(). To account for gene dropout in the single cell data, right aligned moving averages of gene expression were calculated for cells along pseudotime with a window size of 1000 cells. These moving averages were then plotted against pseudotime to show PELI2, IFNB1, and ISG expression over the course of viral infection.

