## Supplementary figures and images for "STING activation of IRF3 is tuned by PELI2 to suppress basal activation and reinforce the anti-viral response"

### Supplemental Figures

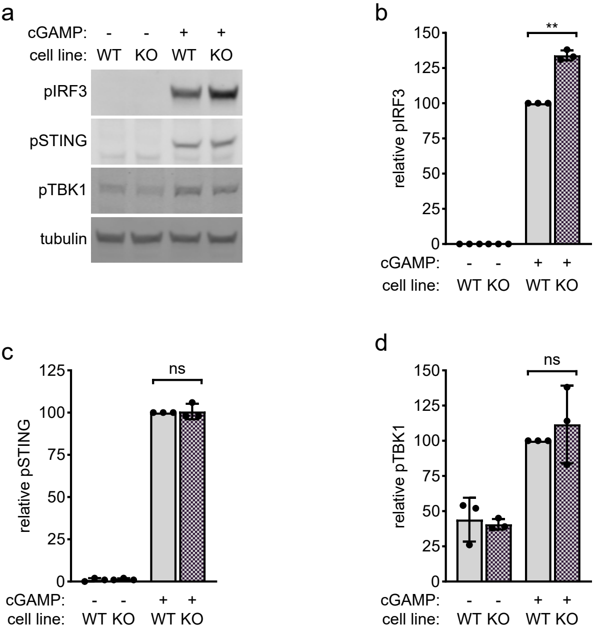

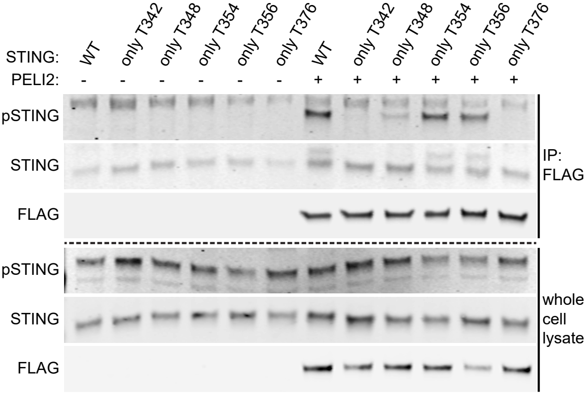

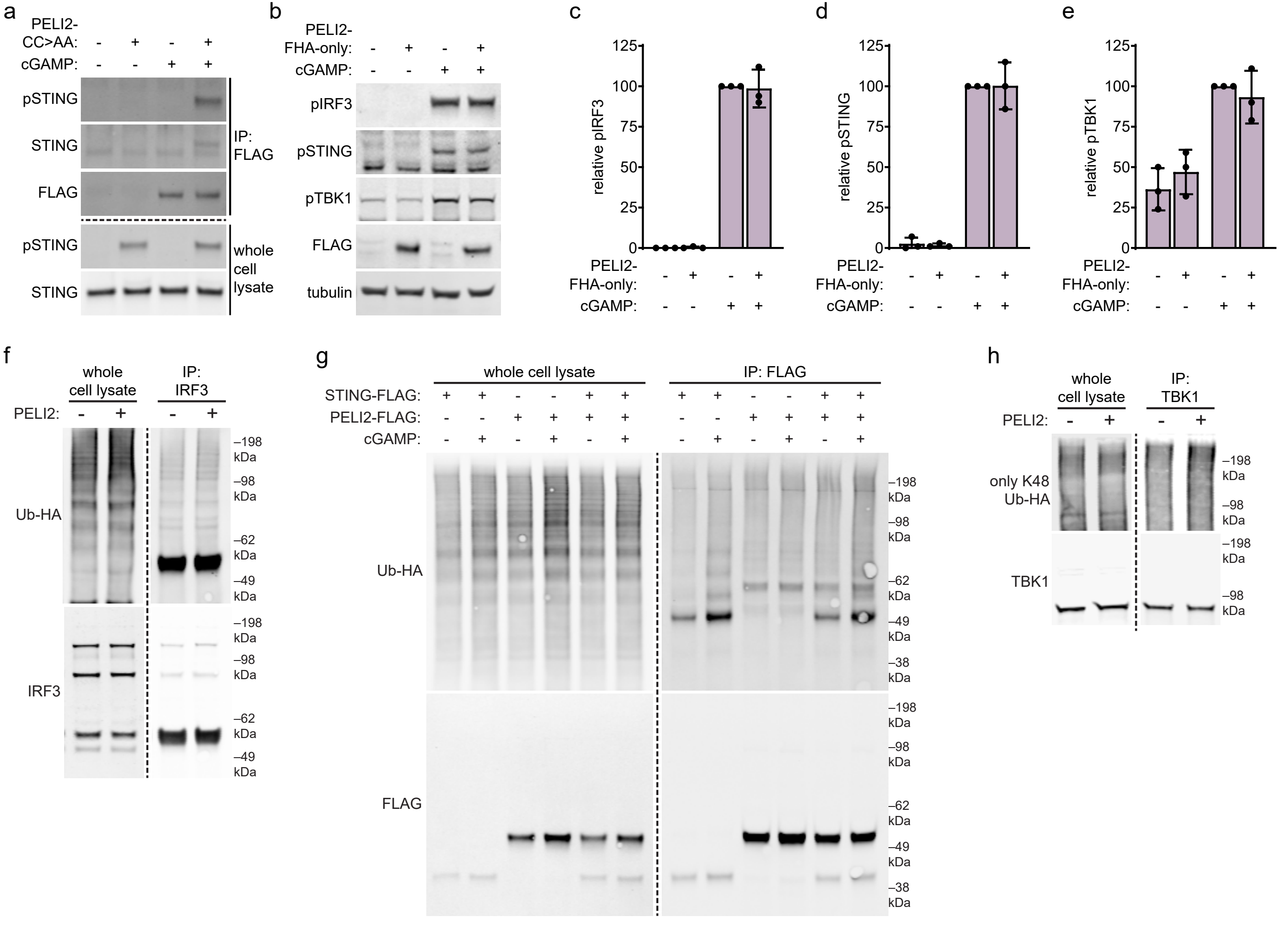

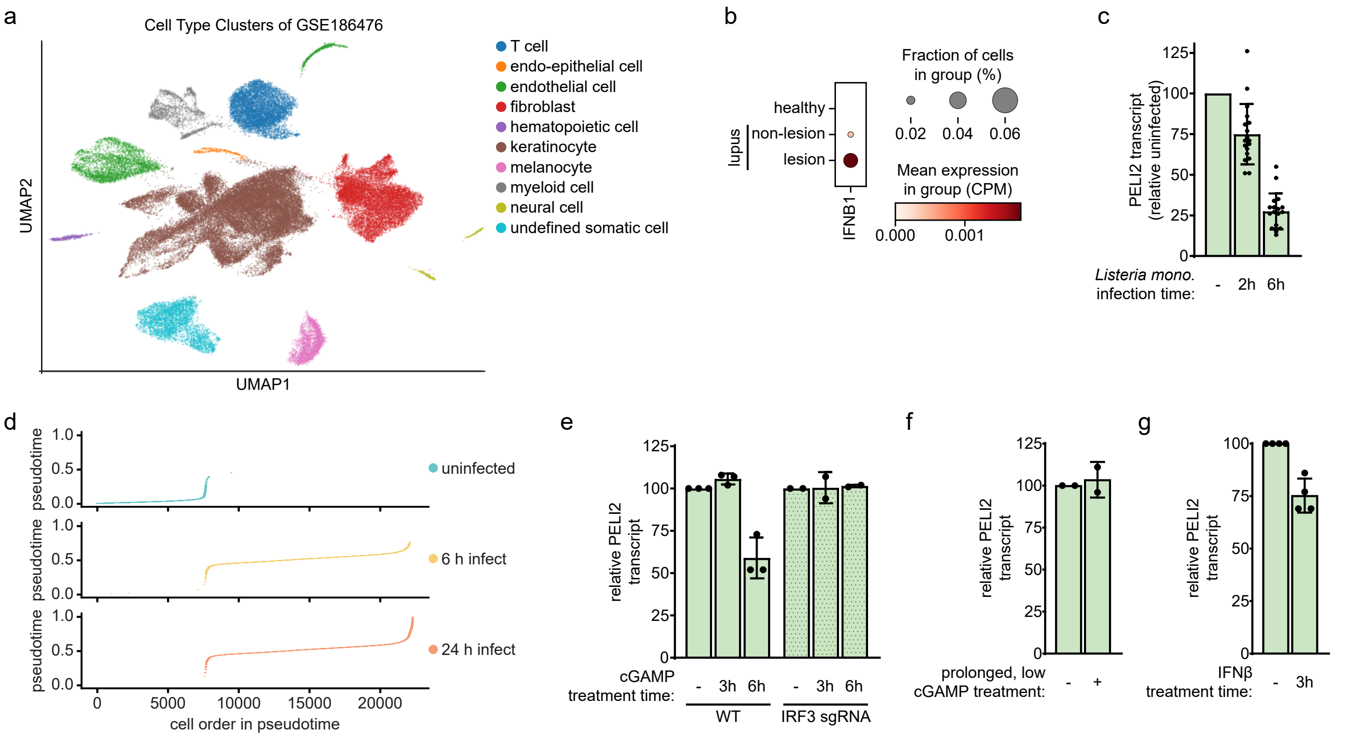

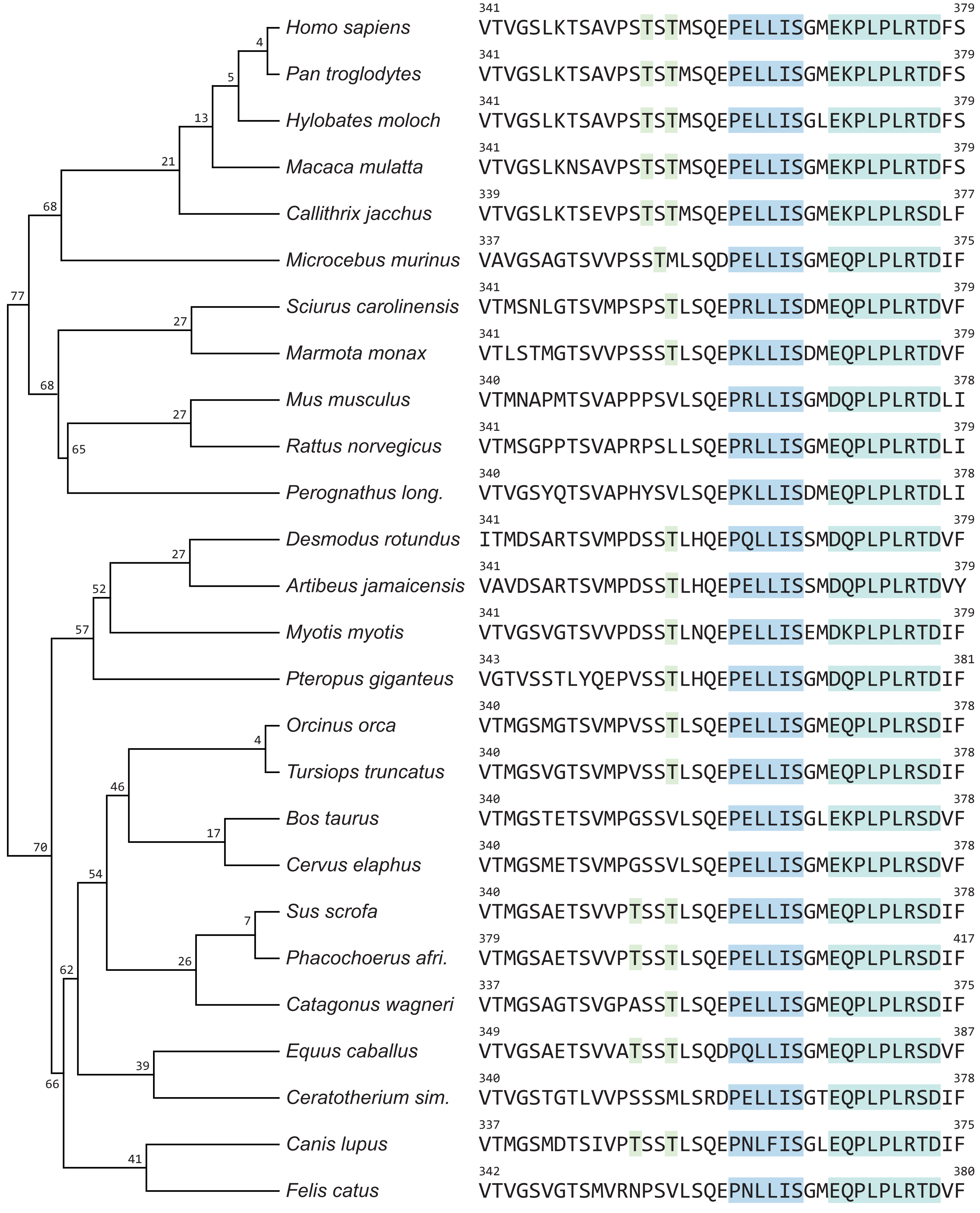
